# Intergenerational effects from spatial and genetic environment predict early-life social network structure

**DOI:** 10.1101/2022.07.19.500597

**Authors:** Victoria R. Franks, Rose Thorogood, Patricia Brekke

## Abstract

Early independence is a crucial stage in the ontogeny of social environments, but it is often challenging to study in the wild. Genetics may structure groups if young animals associate with familiar kin, but association opportunities also develop as a by-product of environmental processes such as spatial resource distribution. The contribution of these alternate factors in initial opportunities for bonding outside direct relatives is difficult to pick apart, despite its importance in shaping later life. However, species where genetics and spatial structure are less closely coupled (for example, via extra-pair mating) provide a natural opportunity to disentangle these effects. We addressed this gap by investigating the contribution of relatedness versus spatiotemporal synchrony (natal nest-box location and fledge timing) to early-life social structure in newly-independent young hihi (*Notiomystis cincta*). We also investigated the contribution of inbreeding in both juveniles and their parents, to individual-level sociality, as this genetic factor has had limited focus in studies of social structure. Using a long-term genetic pedigree, detailed breeding records, and social network data collected across three cohorts, we found that juvenile social associations were predicted by natal nest-box location, irrespective of relatedness between juveniles. Therefore, the physical environment can create initial opportunities for associations to develop once young animals disperse from natal sites. Furthermore, juvenile sociability was predicted by their father’s (but not mother’s) inbreeding, highlighting how genetics may have indirect and intergenerational effects on social behaviour. Overall, social structure in wild animals can emerge early in life if the natal environment determines association opportunities. These patterns may even be pre-determined across generations if breeding and settlement decisions made by parents affect the physical and social environments experienced by their offspring. Ultimately, our study highlights how influences on early life social structure may have important consequences for population dynamics and evolutionary potential.

## INTRODUCTION

Early independence represents a uniquely important, and yet understudied, point in the development of social environments in wild animals. As young leave their parents, they encounter their first opportunities to interact with other individuals outside their immediate family group. Social interactions during this sensitive early-life period have a suite of consequences for later-life behaviour and fitness (Cantor et al., 2021), including sociality (Brandl et al., 2019), senescence (Péron et al., 2010; Turner et al., 2021) and reproductive strategies (Macario et al., 2017; Turner et al., 2021). Consequently, when the structure of the social environment in newly-independent juveniles determines these interaction opportunities they have the potential to delineate boundaries for crucial processes including gene-flow (Sugg et al., 1996) and cultural and social evolution (Cantor et al., 2021; Kuijper and Johnstone, 2019). However, despite a growing body of evidence showing that animals do not interact equally (Sosa et al., 2021; Wey et al., 2008), we have little knowledge of the factors that contribute to social structure in free-living young animals, because juveniles are often mobile, cryptic, and difficult to study in comparison to other life stages which lie either side of the early-independence period.

Genetics are fundamental to the fitness and selection consequences of sociality. Early independence presents an opportunity for juveniles to associate with unrelated individuals for the first time, meaning that genetic relatedness underlying associations may become more variable at this point. In some species, siblings continue to form strong bonds despite the option to disassociate (Archie et al., 2006; Carter et al., 2013; Kurvers et al., 2013), which can enhance social learning or cooperation (Kerth, 2008; Schwab et al., 2008), and improve reproduction, survival, and growth (Chakrabarti et al., 2020; Feh, 1999; Gerlach et al., 2007; Thünken et al., 2016). However, the benefits of kin associations can be context-specific, particularly if social structure predicts later mating decisions (Firth and Sheldon, 2016) and developing associations between genetically dissimilar or unfamiliar non-kin prevents inbreeding (Godfrey et al., 2014; Hirsch et al., 2013; Kurvers et al., 2013; Mourier and Planes, 2021). However, while evidence for kin structuring has been considered in many species, identifying the mechanisms driving this variation remains more challenging. Kin are often inherently more familiar as a product of sharing a nest environment, and other factors such as philopatry may also lead siblings to continue to associate rather than genetic relatedness itself (Leedale et al., 2020).

Alongside influencing association assortment, genetics can also impact on sociality at the individual level via genetically-linked traits: for example, larger individuals may succeed in gaining more associates than smaller conspecifics (Pack et al., 2009). However, genetic effects may also be more indirect. The genotypes of parents can affect traits in their offspring via the environment they create during raising (Kong et al., 2018), known as “ genetic nurturing”. For example, if parents create a poorer or more stressful early-life environment this can in turn affect their offspring’s later-life social strategies (Boogert et al., 2014; Brandl et al., 2019; Farine et al., 2015b). One key aspect of genetics that affects both individual traits and parental care, and may therefore impact on sociality either directly or indirectly, is inbreeding. Firstly, being more inbred alters an individual’s life-history (e.g. condition and secondary sexual characteristics: Bolund et al. (2010)), and behaviour (e.g. personality: Müller and Juškauskas, 2018, but see Herdegen-Radwan, 2019; dispersal strategies: Daniels and Walters, 2000; cooperation: Wells et al., 2020). These traits can affect both quality and quantity of associations: as one example, animals with more reactive personalities have fewer and less stable social network connections (Aplin et al., 2013). Secondly, inbreeding alters parents’ investment strategies in their offspring (Duthie et al., 2016; Pooley et al., 2014; Wells et al., 2020), and can therefore alter early-life raising conditions (Pooley et al., 2014). While inbreeding depression is not always apparent in benign environments (Armbruster and Reed, 2005; Crnokrak and Roff, 1999), the combined challenges represented by parenthood and early-life independence may make the effects of inbreeding on sociality especially apparent in juveniles. However, while this demonstrates how traits and states resulting from inbreeding may be linked to juvenile social behaviour, there have been no studies explicitly testing the relationship between sociality and different generational levels of inbreeding *per se*.

While genetics is often key to the costs and benefits of social living, this component cannot be considered in isolation. The need to consider environmental context when studying animal sociality has become increasingly recognised in recent years (Albery et al., 2021b, 2021a; Evans et al., 2020; Sosa et al., 2021; Spiegel and Pinter-Wollman, 2020; Webber and Vander Wal, 2018). Abiotic factors such as habitat configuration and resource distribution shape how animals interact with a landscape, and in turn affects association opportunities (He et al., 2019). For juveniles, natal site may be a vital environmental factor in shaping their first non-family associations as they disperse away from their natal territory, because juveniles born in territories in close proximity could experience similar habitat effects on their movement and settlement decisions (Bowler and Benton, 2005; Fronhofer et al., 2018). Conversely, if kin disperse away from natal sites to avoid inbreeding (Bowler and Benton, 2005), then we might expect weaker associations between juveniles from similar locations. Ultimately, if natal site acts as a physical factor impacting on social structure, then juvenile association opportunities may be determined even before they are born through breeding and settlement decisions made by their parents (Ilany and Akçay, 2016).

The likelihood that juveniles will co-occur in space also varies temporally (Pasquaretta et al., 2021; Psorakis et al., 2015; Whitehead and Dufault, 1999). During the breeding season, variation in resources at global and local scales can create variation the timing of mating, gestation, and eventually when young become independent (Ringsby et al., 2002). Thus, if juveniles become independent at the same time, they may be more likely to encounter each other, particularly in combination with spatial effects or if there are also age-associated preferences in habitat use (Ferrer and Penteriani, 2003) or social tendencies (Turner et al., 2017). However, explicitly testing for combined spatiotemporal effects is often largely overlooked in studies of animal sociality (He et al., 2019; Sosa et al., 2021; Wey et al., 2015). The effects of relatedness, environment, and timing are often confounded because siblings from the same clutch or litter are inherently born in the same place at the same time, and so understanding the individual contributions of each component to natural social structure is often impossible. Furthermore, teasing apart these effects also requires a high level of detailed breeding and genetic data, which are rarely available from wild animal populations (Clutton-Brock and Sheldon, 2010). As a result, multivariate analysis of complex combined factors contributing to animal social structure (such as space, time, and relatedness) is challenging, and few studies have begun to tackle these areas in detail (Sosa et al., 2021; Wolf and Trillmich, 2008).

While disentangling the contributions of genetics, space, and time to early-life social structure is challenging in wild populations, some species do provide a natural opportunity to unpick these relationships. The hihi (*Notiomystis cincta*), a threatened New Zealand passerine, has one of the highest known rates of avian extra-pair paternity (mean frequency of EPP in broods reported as 0.68±0.012 by Brekke et al. 2013), meaning that not all nest-mates are closely related and close kin are not only within nests. In this study, we combined this natural phenomenon with a wealth of detailed genetic and breeding data available on our focal study population (Tiritiri Matangi Island, New Zealand) to provide a rare insight into the factors contributing to early-life social environment structure in wild animals. On Tiritiri Matangi, hihi nest in boxes that we provide, which has enabled all breeding attempts to be monitored and recorded since the population was established in 1995. Blood sampling of all individuals has occurred since 2005 for genetic analysis, creating a long-term pedigree with derived inbreeding and relatedness metrics (Brekke et al. 2015). Finally, once juvenile hihi (i.e., offspring from the current breeding season) disperse from their natal nests, they congregate at 2-3 spatially separate sites for approximately four months each year (Franks et al., 2020b). Sociality in young hihi has important influences on behaviour (Franks et al., 2020c) and survival (Franks et al., 2020a), but social structure had not been examined at a detailed genetic or spatiotemporal level prior to this study. We used social network analysis to analyse group- and individual-level social structure, which has become a widely-applicable tool to quantify non-uniform associations between animals (for review, see Krause et al., 2009; Sosa et al., 2021; Wey et al., 2008). First, we investigated if associations between juvenile hihi in each group site were stronger if birds were more related and had fledged from nest-boxes that were closer together in time and/or space. Secondly, we tested the contribution of genetic factors to individual-level sociality, by analysing whether juveniles’ number of associates depended on their own or their parent’s inbreeding.

## METHODS

### Study population

Our study was conducted over three years (2015 – 2017) on Tiritiri Matangi Island (New Zealand, 36°36’00” S 174°53’21” E). The study site is a 2.5 km^2^ island covered in a mixture of native subtropical rainforest and more open grassland. Hihi on Tiritiri Matangi are monitored intensively during the breeding season, which is typically from early September to February. Each year, parent birds hold a territory and raise altricial young (up to three clutches) in the nest-boxes we provide. Both parents contribute to feeding offspring once hatched (Castro et al., 1996). Nest-boxes are checked following an established protocol (Ewen et al., 2018) which allows us to record the outcomes of all nesting attempts, and identify the dam and social male caring for each brood. Blood samples are taken from all nestlings at 21 days old (hatch day = day 0) to establish genetic sires and maintain a genetically resolved pedigree. Nestlings are also ringed with unique combination of coloured leg rings at this age. Young hihi fledge at around 28 days old, and after approximately two weeks of post-fledging dependency they disperse from their natal territory to congregate at 2-3 spatially separate sites in forested gullies for approximately four months (Figure 1; for more details see also Franks et al., 2020b).

**Figure 1.**
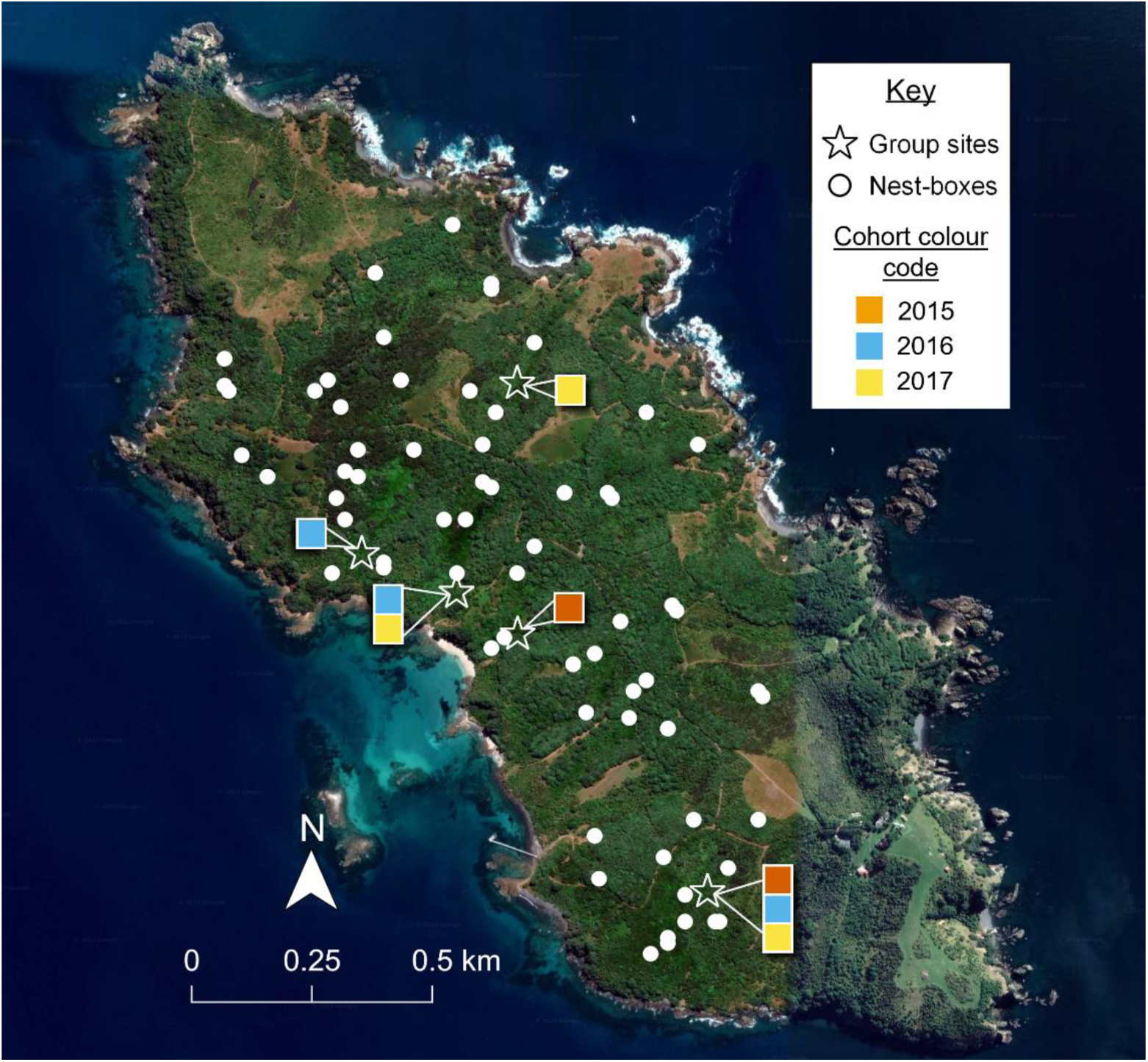
Map showing the location of juvenile hihi grouping sites (white stars: 2015 = red; 2016 = blue; 2017 = yellow) and nest-boxes (white circles) included in this study. Map data: Google, Maxar Technologies, TerraMetrics, CNES/Airbus (2022).

During the three years of our study, the hihi population varied between 180 and 270 individuals, with similar proportions of juveniles and adults (second year or older) each year (McCready and Ewen, 2017, 2016; Smith and Ewen, 2015). The first year of our study (2015) was a poorer breeding season than 2016 and 2017 (2015: 89 fledglings; 2016: 132 fledglings; 2017: 151 fledglings), which we accounted for in analyses. There were two juvenile group sites in the 2015 season, and three group sites in both 2016 and 2017 (Figure 1; see also Franks et al., 2020b).

### Ethical note

Ethical approval for the observations of juvenile groups was issued through the Zoological Society of London Ethics Committee (UK). All yearly monitoring of breeding populations (including nest-box monitoring, blood sampling and colour-ring application) followed an established protocol conducted under a New Zealand Department of Conservation Research Permit (authorisation number: 44300-FAU).

### Social network data

Each year from mid-January to April, we observed juvenile hihi as they congregated at group sites (15 observation sessions per site in 2015, 25 sessions per site in 2016 and 2017). Each observation session lasted one hour, which was sub-divided into 30-second time blocks (120 blocks). In every 30 second block, we recorded the coloured leg-ring combinations of all hihi perched within a 15-metre radius of the observer. Any bird present across multiple blocks was re-recorded at the start of each block, so we could determine continued presence to the nearest 30 seconds. This system was used as the most fine-scale timing possible to capture changes in presence of small forest passerines while also being long enough to allow for identification and recording of individuals. All observations were made with binoculars (Zeiss Conquest® HD 8×42) by one observer (VRF). Observations were made from the same point in each group site for each observation session, and sessions were distributed evenly across the three months each year.

We constructed an undirected weighted social network for each cohort using the R package *asnipe* (Farine, 2013), which defined associations based on spatiotemporal proximity in our time-stamped observations of juveniles; thus, we assumed individuals seen often in the same location at similar times were more familiar (gambit of the group approach (Whitehead, 2008)). Each association value represented how frequently each pair of juveniles (dyad) associated (from 0, never seen together to 1, always seen together). Details on each cohort, including total numbers of individuals included in each network and number of network associations, are summarised in Supplementary Table 1.

### Breeding data

#### Relatedness and inbreeding coefficients

A pedigree has been maintained in this population since 1995, with paternal resolution available from 2005 when molecular markers became available (see Brekke et al., 2015). Maternity is determined during nest monitoring by identifying the female that broods each clutch and cares for the chicks. Paternity is determined using genetic information from the blood samples obtained from hihi nestlings. Genomic DNA is extracted and screened using 18 neutral microsatellite loci (15 species-specific and three isolated from other passerines) that are widely distributed across the genome (complete methods in Brekke et al. (2013)).

Using the pedigree, we calculated relatedness (*r*) between all dyads in each social network using the R package *nadiv* (Wolak, 2012). Relatedness ranged from 0 (no common ancestors), to approximately 0.5 (full siblings), although exact relatedness values could be slightly higher due to historical inbreeding (maximum *r* = 0.7) Pedigree-derived inbreeding coefficients (*F*, the expected proportion of an individual’s genome that is identical by descent) were also calculated using the *nadiv* package. Relatedness among individuals and inbreeding values were both estimated using a minimum of six known ancestors (parents and all grandparents) to reduce metric bias due to pedigree depth. Any birds which did not have known parents and grandparents were removed from further analyses.

#### Natal nest-box location and fledging synchrony

The latitude and longitude for all nest-boxes (distributed across the 2.5km^2^ island) were recorded as part of the ongoing monitoring of this population (Figure 1). We defined the distance (in km) between the nest-boxes of every juvenile dyad in each social network using the *geodist* R package (Padgham and Sumner, 2021), which calculated geodesic distance based on shortest path length between coordinates. The resulting matrix of distances provided a spatial layout of proximities between the natal nest-boxes of each pair of juveniles (from 0 – 1.55km).

Finally, a fledging date was recorded for each clutch as part of monitoring of all breeding attempts on Tiritiri Matangi: this was the date the last chick left the nest (i.e., active nest-boxes near to fledging were checked daily until they were found empty). Therefore, for our analyses each juvenile within a clutch was assigned the same fledge date. We then calculated the number of days between fledge dates for every pair of juveniles in each cohort’s network to give a “ fledge synchrony” value for every dyad.

### Dyadic analysis of pairwise associations

To investigate how relatedness, distance between natal nest boxes, and fledge synchrony predicted associations in dyads of juvenile hihi, we fitted Bayesian logistic mixed-effects model using a Markov Chain Monte Carlo (MCMC) framework. We used the *MCMCglmm* package (Hadfield, 2010), because its multi-membership modelling capabilities allowed us to account for each individual appearing interchangeably in a dyad (i.e., a hihi could be individual A or individual B in each association in our undirected networks). For every model, we used non-informative priors and adjusted burn-in periods, iterations, and thin intervals to ensure minimum autocorrelation and good convergence (determined in post-hoc diagnostic checks) (Hadfield, 2010). We inspected models for goodness-of-fit using the Deviance Information Criterion (DIC). To determine the predictors that best explained our data, we removed non-significant terms from the global model to obtain the most parsimonious model which had good fit (low DIC) (Rushmore et al., 2013). In all analyses, we removed dyads that included any individuals that had been seen fewer than five times, because initial exploration for outlier association strengths in our data indicated that dyads observed fewer than this amount had the least reliable network measures.

Many juveniles never associated in our networks (Supplementary Table 1) and different mechanisms could affect opportunity to associate versus strength of association when they did occur. Thus, we used a stepwise approach to investigate juveniles’ (a) likelihood of association (binary response: 1 = associated, 0 = never associated), then (b) association strength. We first compared association likelihoods between nest-mates and non-nest-mates, to understand whether sharing a nest was important for juvenile social structure. In a binary MCMC model, our main predictor was whether dyads had fledged from the same nest (categorical value where same nest = 1, different nests = 0). Next, we considered association likelihood in juveniles from different nests, who had variable nest-box proximities and fledge dates and thus provided an opportunity to explore the effects of spatiotemporal variation on broad-scale network structure. Again, we used a binary response (1 = associated, 0 = never associated); here, the main predictors were distance between natal nest boxes, fledge synchrony, and additionally relatedness between dyads to understand whether genetic sibling recognition influenced social structure. Finally, we accounted for potential biases in association metrics (Franks et al., 2021): we specified the fewest number of observations for each pair as a predictor (i.e. if individual A was seen 10 times and individual B 20 times, minimum observations = 10), and included cohort year as a random effect as association opportunities may have varied among cohorts.

We then considered (b), variation in association strengths in dyads that had associated at least once. For all following models using association strength, we z-transformed the values so that they were comparable among the three cohorts in our study. Nest-mates all fledged on the same day and from the same location so had values of 0 for nest-box location and fledge synchrony, but their relatedness could vary from ≥0.25 due to EPP; meanwhile, non-nestmates fledged on different days from different locations, and had relatedness of ≤0.25. Therefore, we investigated association patterns in same-nest and different-nest juveniles separately. To analyse how relatedness predicted association strengths between nest-mates. our main predictor in our MCMC model was whether nestmate dyads were half or full siblings, which we determined from pedigree relatedness values as they showed a clear bimodal distribution (half siblings: *r* < 0.5; full siblings: *r* > 0.5; Supplementary Figure 1). We also included a predictor that quantified whether juveniles originated from nests that contained only half-siblings, full-siblings, or a mix of both sibling types: even though there were similar numbers of dyads from each nest type which provided equal opportunities for association (Supplementary Table 2) this parameter allowed us to quantify if association strengths were consistent even when juveniles had the opportunity to associate with both sibling types. Finally, in a separate model we explored association strengths in juveniles from different nests, where relatedness, distance between natal nest boxes, and fledge synchrony all varied. Again, the response variable was z-transformed association values, and our main predictors were relatedness, distance between natal nest-boxes, and fledging synchrony. Here, relatedness values could not easily be categorised into full-and half-siblings, so we used this variable as a continuous predictor. Again, to account for potential biases in association values we also included the fewest number of observations for each pair in both models analysing association strengths, and a random effect for year to examine overall patterns across the different cohorts.

### Inbreeding and sociality analysis

To examine whether sociality was predicted by both an individual’s own level of inbreeding and the inbreeding of its social parents, we calculated the degree strength score for each juvenile, which quantifies the number and strength of associations and is thus a measure of centrality in the network (Krause et al., 2015). We then used a Generalised Linear Mixed Effects model (GLMM) with each juvenile’s degree strength as the response variable, which we z-transformed so we could examine general trends across the years. Our main predictors were the juvenile inbreeding coefficients, mother’s coefficients, and father’s coefficients; all inbreeding coefficients were z-scored. These variables were checked for collinearity prior to inclusion in the model, but no issues were found. We also included an interaction between the father’s inbreeding coefficient and whether they were the full genetic father, or the social father only (not a direct parent but providing care during chick-rearing). We specified a random effect of nest-box identity, because multiple fledglings originated from the same nests and shared the same mothers and fathers.

Analyses using social network metrics violate the assumptions of many statistical tests due to non-independence of data, which can lead to inflated Type-I and Type-II error rates (Farine and Whitehead, 2015). Furthermore, biases can be introduced from sampling effort or spatiotemporal variation, and need to be accounted for to ensure that results are valid. Following recent advances in network analysis (for discussion, see Farine and Carter, 2022; Franks et al., 2021; Weiss et al., 2021) we used the double permutation procedure proposed by Farine and Carter (2022) to calculate the significance of our effects. Here, data stream permutations first calculate any potential deviation in network metrics due to unwanted biases (e.g. spatial or sampling effects). The corrected metrics are then used to investigate the effect of each parameter of interest, with node permutations used to calculate statistical significance. As well as using double permutations to reduce any effects of observation bias among individuals, we also removed individuals observed fewer than 20 times as these individuals had the least reliable degree strengths based on initial examination of the data.

## RESULTS

### Effect of relatedness, fledging times, and nest-box proximity on social structure

#### Association likelihoods

We found that the social network structure of juvenile hihi was predicted by spatial characteristics of the natal environment. Associations were the most likely to form between juvenile hihi that had been nest-mates: sharing a natal nest-box was included in our final binary model exploring association likelihood across all juveniles (*N* dyads = 11184, *N* birds = 171, Table 1a, Figure 2). Additionally, when we considered juveniles from different nests (*N* dyads = 10924, *N* birds = 171, nest-box proximity was included the model that best explained association likelihood (Table 1b). Therefore, non-nestmates tended to form associations if they fledged from boxes located more closely together (Table 1b, Figure 3). By contrast, there was no evidence that association likelihood changed if juveniles had fledged at similar times, or if the dyad was more closely related, as neither fledge synchrony, relatedness nor any interaction between our three main predictors were included in the final model. Finally, there was also a minimal effect of number of observations on network structure, because minimum number of observations per dyad was included in both final models investigating association likelihood but the actual effect sizes were small (Table 1).

**Table 1.**
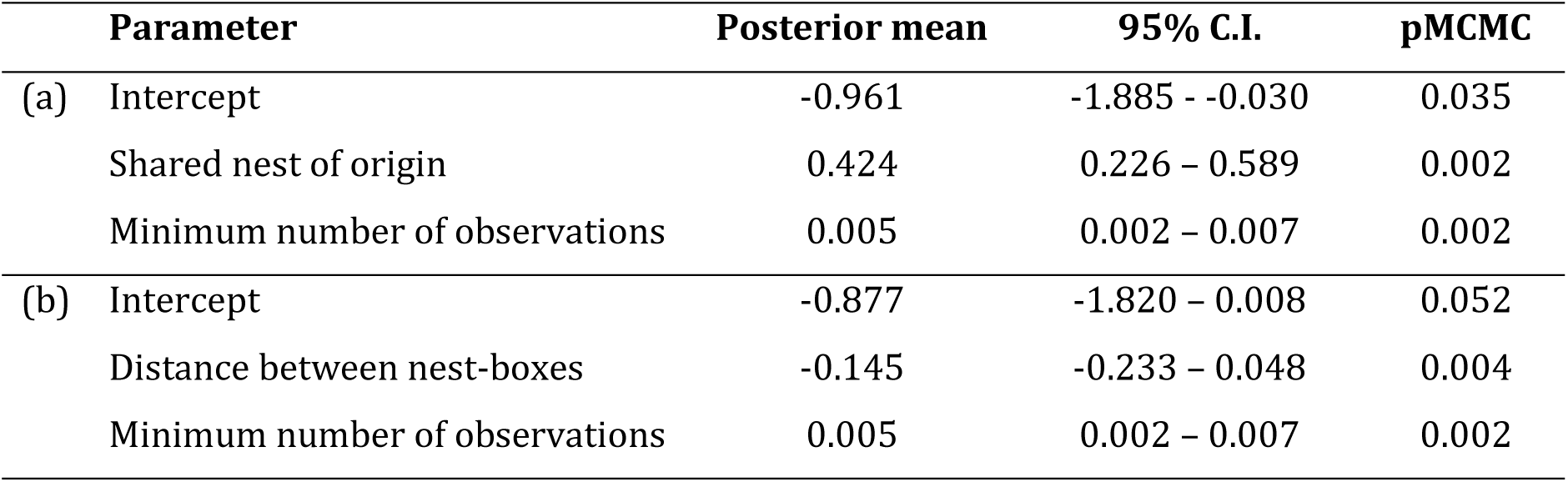
Predictors included in the final models analysing binary dyadic association likelihood in three cohorts of juvenile hihi, considering (a) associations among all juveniles, and (b) associations within non-nestmates only. Posterior means, 95% credible intervals, and P-values were calculated with a Bayesian logistic mixed-effect approach using Markov chain Monte Carlo sampling. Non-significant parameters were removed from the final models.

|  | Parameter | Posterior mean | 95% C.I. | pMCMC |
| --- | --- | --- | --- | --- |
| (a) | Intercept | -0.961 | -1.885 - -0.030 | 0.035 |
|  | Shared nest of origin | 0.424 | 0.226 – 0.589 | 0.002 |
|  | Minimum number of observations | 0.005 | 0.002 – 0.007 | 0.002 |
| (b) | Intercept | -0.877 | -1.820 – 0.008 | 0.052 |
|  | Distance between nest-boxes | -0.145 | -0.233 – 0.048 | 0.004 |
|  | Minimum number of observations | 0.005 | 0.002 – 0.007 | 0.002 |

**Figure 2.**
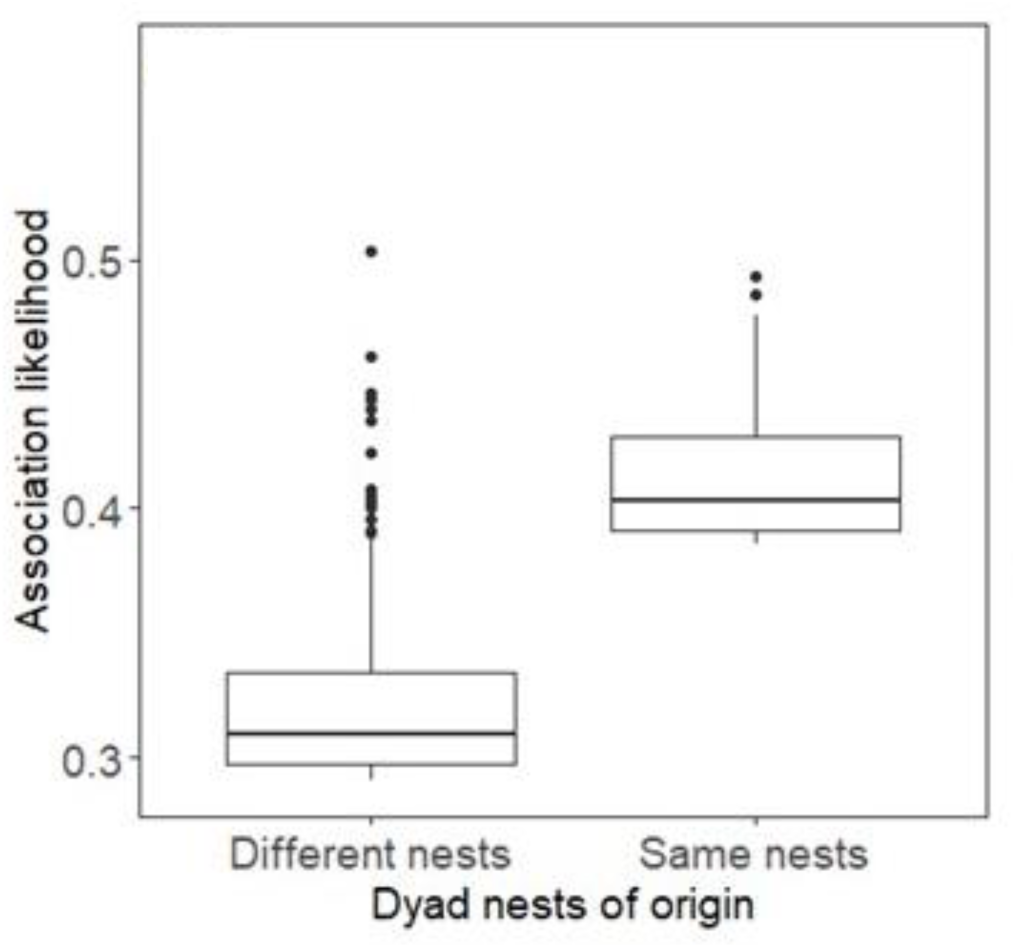
Likelihood of association between dyads of juveniles originating from different nests, and the same nests. Values are predicted from the final model exploring association likelihood across all juveniles (Table 1a)

**Figure 3.**
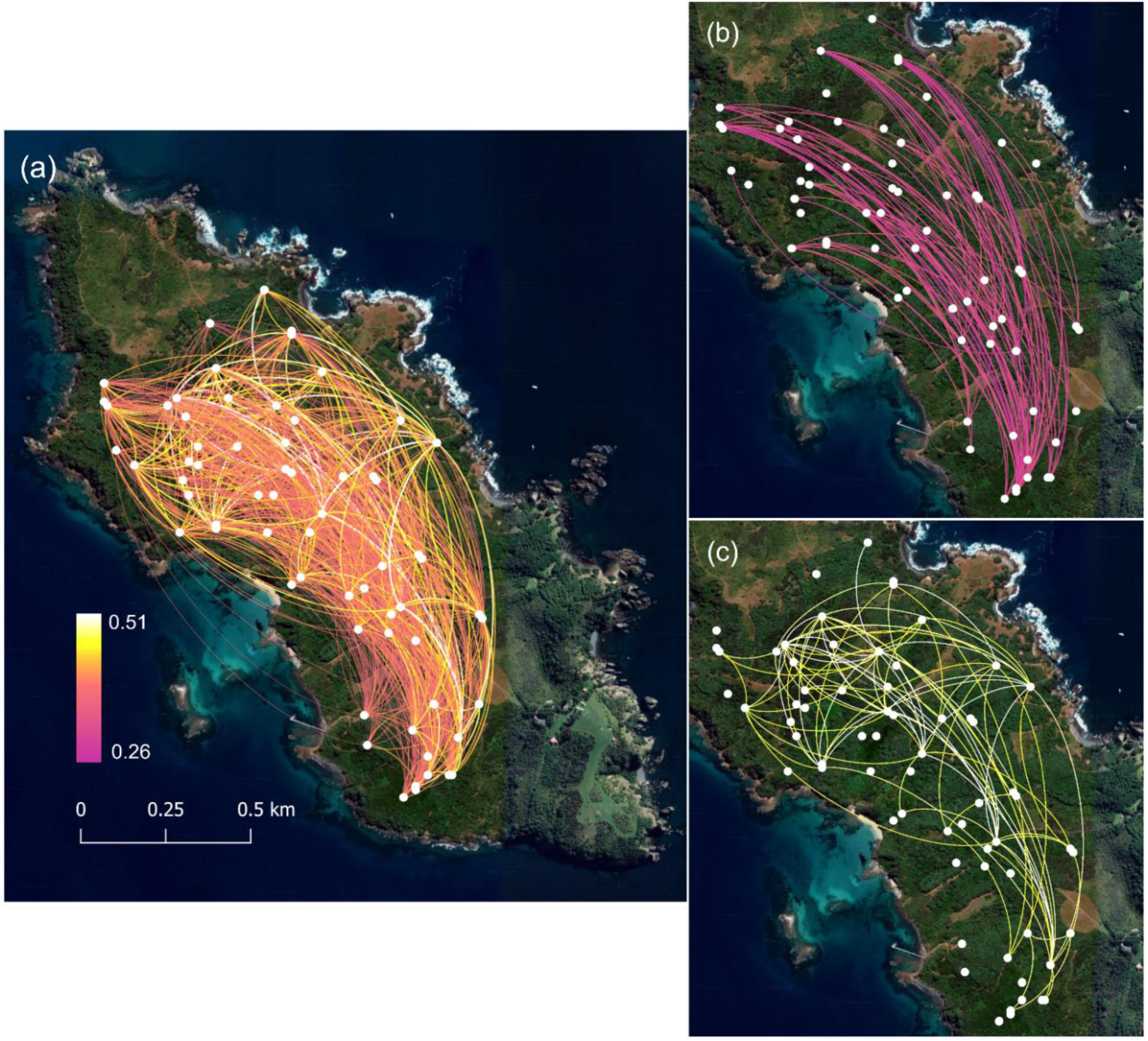
Likelihood of associations forming between juveniles from different nest-boxes, as predicted by the model in Table 1b; (a) all predicted associations, (b) the 100 least likely and (c) the 100 most likely associations. All map data: Google, Maxar Technologies, TerraMetrics, CNES/Airbus (2022).

**Figure 3.**
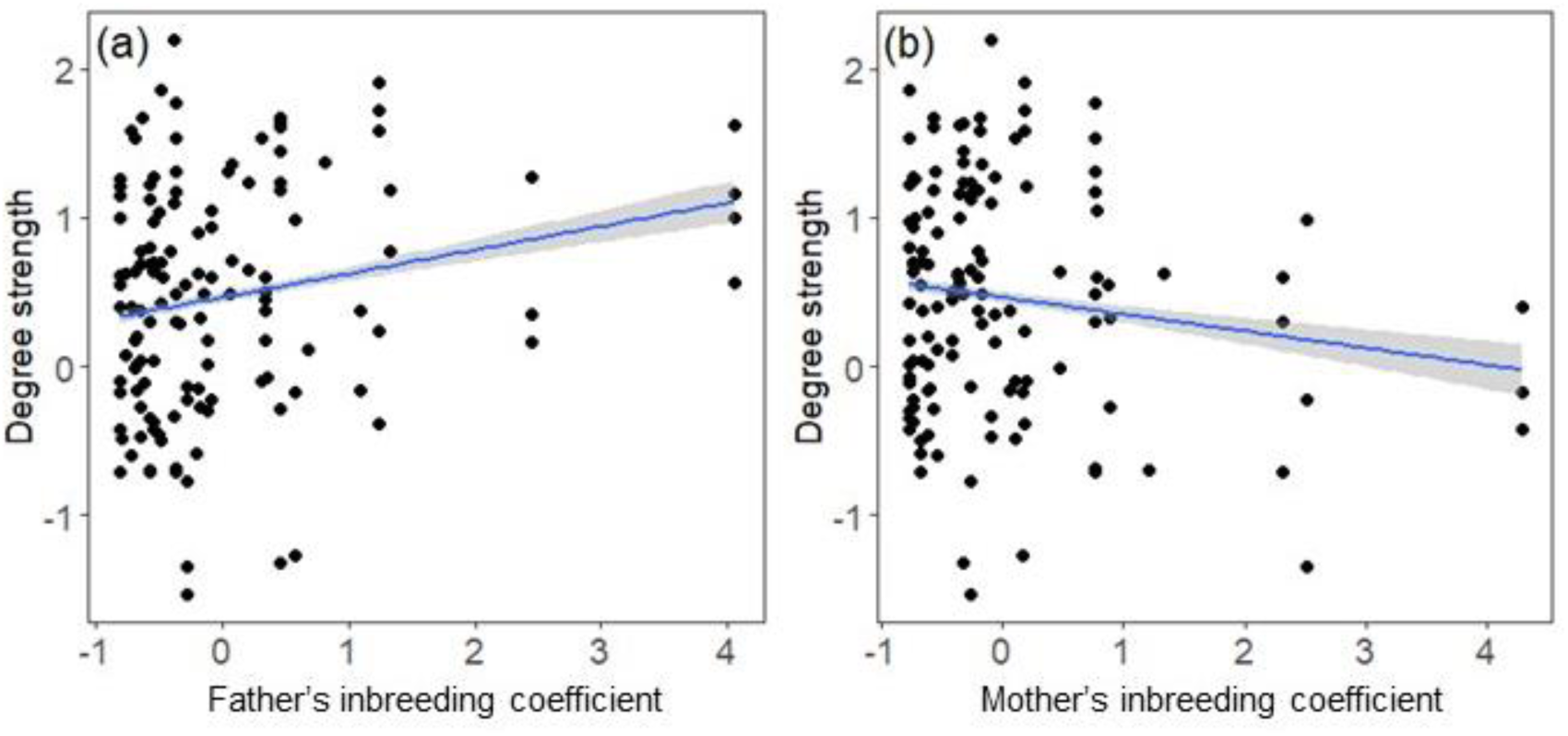
Relationship between the degree strength of each juvenile and their (a) father’s, and (b) mother’s inbreeding coefficient. Degree and inbreeding coefficients are z-transformed. Line of best fit (blue line) with 95% C.I. calculated from model in Table 3.

#### Association strengths

In contrast to our results from likelihood of association, we did not find evidence that our three main parameters (nest-box proximity, fledge synchrony, and relatedness) predicted association strength, as these parameters were not included in the final models for either nestmates or non-nestmates (nestmates: *N* dyads = 2924, *N* birds = 167; non-nestmates: *N* dyads = 2842, *N* birds = 167). In nest-mates, no parameter significantly explained association strength, and the only parameter in the final model for non-nestmates was minimum number of observations, which had a negligible effect on associations (Table 2).

**Table 2.** Predictors included in the final model analysing dyadic association strengths in juvenile hihi, for non-nestmates. Posterior mean, 95% credible interval, and P-values are calculated from a Bayesian logistic mixed-effect model using Markov chain Monte Carlo sampling. Non-significant parameters were removed from the final model.

| Parameter | Posterior mean | 95% C.I. | pMCMC |
| --- | --- | --- | --- |
| Intercept | 0.001 | -0.171 – 0.174 | 0.994 |
| Minimum number of observations | 0.004 | 0.002 – 0.006 | 0.003 |

### Effect of early-life inbreeding environment on sociality

An individual’s own level of inbreeding did not predict its number of network associates (degree strength) (Table 3; *N* = 111 birds). However, a juvenile’s sociability was significantly predicted by its social father’s inbreeding. We found that juveniles whose fathers were more inbred (higher inbreeding coefficient) had higher degree strength and were thus more social (Table 3, Figure 3a), independent of whether the father was genetically related to the juvenile or was only their social care-provider (interaction between father’s inbreeding coefficient and genetic status non-significant, Table 3). This effect remained consistent even when we removed two fathers who had very high inbreeding coefficients, so these individuals were not driving the relationship between father’s inbreeding and juvenile sociability (Supplementary Table 3a; Supplementary Figure 2a). While there was a trend for juveniles with more inbred mothers to be less social (Table 3, Figure 3b), this was not statistically significant in either the full dataset or with two mothers with extreme inbreeding coefficients removed (Supplementary Table 3b; Supplementary Figure 2b).

**Table 3.** Results of linear mixed effect model analysing the effect of inbreeding on individual sociality (degree strength) in juvenile hihi. P*double permutation* is the P-vale resulting from the double permutation procedure, which controlled for number of observations. Significant effects (at *P* < 0.05) highlighted in bold.

| Parameter | Est. | S.E. | t-value | $P_{\text{double permutation}}$ |
| --- | --- | --- | --- | --- |
| Intercept | 0.569 | 0.097 | 4.823 |  |
| Juvenile inbreeding coefficient (z-score) | 0.026 | 0.063 | 0.413 | 0.338 |
| Mother's inbreeding coefficient (z-score) | -0.108 | 0.075 | -1.452 | 0.078 |
| **Father's inbreeding coefficient (z-score)** | **0.257** | **0.124** | **2.074** | **0.017** |
| Genetic status of father (within-/extrapair) | 0.015 | 0.139 | 0.110 | 0.465 |
| Father's inbreeding coefficient*genetic status of father (within-pair/extra-pair) | -0.180 | 0.143 | -1.254 | 0.100 |

## DISCUSSION

Understanding the structure of the early-life social environment is crucial, because interactions in this period have the potential to determine behaviours through to adulthood (Slagsvold and Wiebe, 2011) and influence key population processes such as survival and reproduction (Nuñez et al., 2015; Slagsvold and Wiebe, 2011). However, we often have limited opportunity to understand how different socioecological components impact on associations once juveniles leave their parents to determine the types of interactions available. In this study, we investigated the contribution of spatial, temporal, and genetic factors to early-life social structure in groups of juvenile hihi. Across three cohorts of juveniles, we found that individuals from the same nest were most likely to form associations. Similarly, juveniles from different nests showed a tendency to associate when their nest-boxes had been closer together. However, we found no evidence for effects of nest-box proximity, synchrony in fledging timing, or relatedness on finer-scale association strengths. Finally, each juvenile’s sociability (degree strength) was not predicted by their own inbreeding, but instead related to their parents’: in particular, juveniles were more social if their fathers were more inbred. Overall, these results highlight the dual importance of ecological and genetic components from early life in determining social interactions between juvenile animals, and demonstrate how these different factors may act at different levels within populations to create an emergent social structure from a young age.

The physical structure of the environment, such as resource distribution and spatial configuration, shapes both individual and collective behavioural decisions which in turn determines when and where animals co-exist in space and time (Mbizah et al., 2020; Pasquaretta et al., 2021; Sosa et al., 2021; Strandburg-Peshkin et al., 2017). In our study, we extend knowledge on the importance of habitat on social structure by highlighting how the physical natal environment contributes to the first opportunities that young animals have to associate once they are independent. In hihi, the location of nest boxes predicted early-life association likelihood: juveniles from the same nest were most likely to form associations. Association likelihood then declined as distance between nest-boxes increases, although this effect was small which may be because the size of our study site (2.5km^2^) limited the extent that associations could differentiate. Nevertheless, the combined evidence within and across nests highlights how an animal’s physical environment can predict the basis of social structure when some animals to associate, but not others (He et al., 2019). For juveniles, natal location may create association opportunities through one major process at this life stage: dispersal is a key event for determining the future distribution of juveniles in a population and can be shaped by the surrounding configuration of resources including food and suitable habitat (Kaemingk et al., 2019; Messier et al., 2012; Paradis et al., 1998). Thus, juveniles from similar natal locations may share similar dispersal patterns, making natal location important in determining the opportunities for the ontogeny of social associations at the initial point of early-life independence.

Relatedness and familiarity are often inherently linked within siblings due to their shared raising environment and are difficult to tease apart (Leedale et al., 2020): even when studies have found evidence that siblings preferentially associate over unrelated individuals, they rarely separate genetic relatedness from more simple cue-based familiarity (e.g. Bonadonna and Sanz-Aguilar, 2012; Kurvers et al., 2013). However, in our cohorts of wild hihi, we also had an opportunity to separate genetic effects on associations from ontogenetic familiarity because relatedness varied within and across nests due to EPP. Overall, there was a lack of evidence for genetic relatedness underlying dyadic association patterns, because more closely related individuals were not more likely to be connected in our networks. This supports a recent review that concluded genetic cues to kinship are rare in birds overall, while familiarity from learned or environmental cues offer a more parsimonious explanation for associations between kin in most contexts (Leedale et al., 2020). Alongside the finding that associations were most likely between nest-mates, this highlights the importance of natal environment for association opportunities, independent of any link with relatedness between associating individuals.

Social structure may be further mediated by traits and/or states that affect the number and strength of associations individuals have (Croft et al., 2009; Farine et al., 2015a; Gartland et al., 2021; Pike et al., 2008). However, in our study we found no evidence that a juvenile hihi’s own extent of inbreeding affected its social behaviour. Instead, their individual sociability was predicted by their fathers’ extent of inbreeding, irrespective of whether this individual was their genetic or social father, which indicates there may also be more indirect link between inbreeding and social behaviour acting across generations. Inbreeding causes parents to alter how they invest in their young, affecting resource allocation (Duthie et al., 2016) and the extent of care provided to young both before and after birth (Pooley et al., 2014; Wells et al., 2020). These conditions created by parents affect the overall raising environment experienced by their offspring, which has been shown to play a crucial role in determining the later-life social strategy of juveniles via stress-linked effects (Boogert et al., 2014; Farine, Spencer, et al., 2015), potentially through acting as a cue for environmental conditions (English et al., 2015). Previous studies have also demonstrated how *when* juveniles experience particular environmental conditions also impacts on the magnitude and directionality of effects on their social behaviour (Boogert et al., 2013). This may explain why we only found evidence for an effect from fathers, and not mothers, if variation in parental roles between the sexes mean that males and females contribute to care in different ways and at different times to exert differential effects on the raising conditions of their offspring (Buitron, 1988; Wesolowski, 1994). Overall, this study provides the first evidence for consequences from indirect and intergenerational genetic effects on the sociability of young animals. To further investigate this relationship, the mechanisms underlying the links between parent inbreeding and offspring sociability now need to be examined more closely, alongside direct measures of parental social behaviour and investment at different stages of offspring care.

Our results support recent developments in understanding how complex social structure emerges from effects acting at a multitude of levels (Cantor et al., 2021), whereby the physical environment influence the likelihood of two animals ever forming an association (Strandburg-Peshkin et al., 2017), but individuals also exhibit their own social tendencies as a product of their traits and experiences which mediates associations once formed (Farine et al., 2015a). Here, we show that overall social structure in wild animals may emerge very early in life if the natal environment determines associations at both the individual and dyadic level. Furthermore, these patterns may even be pre-determined across generations if breeding and settlement decisions made by parents then determine the physical and social environments experienced by their offspring (Ilany and Akçay, 2016). For small populations in particular, the resulting social structure may have consequences for their evolutionary potential depending on individual mixing, particularly if associations continue to determine reproductive decisions across generations (Firth and Sheldon, 2016). However, such conditions may also create environments that promote alternate reproductive strategies in order to buffer the risk of inbreeding, including extra-pair mating that allows genetic relatedness to be more independent of spatial proximity (as seen in our hihi population), or postcopulatory mechanisms of inbreeding avoidance (Brekke et al., 2012). While the consequences remain to be tested explicitly, the potential implications highlight how influences on early life social structure may scale up to fundamentally determine population dynamics and evolutionary potential via the social environment.

## ACKNOWLEDGEMENTS

VF was funded by a research grant from the Association for the Study of Animal Behaviour. We would like to thank Matthew Silk for providing invaluable advice on social network analysis, and Mhairi McCready and Donal Smith for monitoring the hihi population between 2015-2017. We are grateful to the New Zealand Department of Conservation and Supporters of Tiritiri Matangi for their permission to conduct this research.

