## Supplementary information for "Intergenerational effects from spatial and genetic environment predict early-life social network structure"

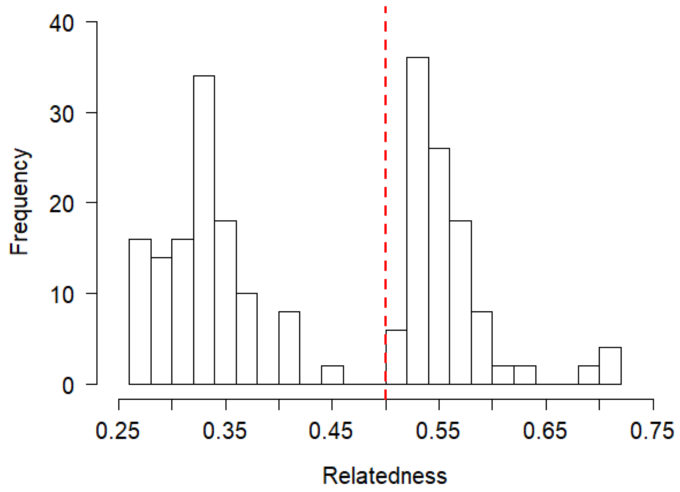


**Supplementary Figure 1.** Distribution of relatedness values for juvenile dyads from the same nests. Red dashed line shows split in relatedness values for half-siblings (*r* < 0.5) and full siblings *r* > 0.5).

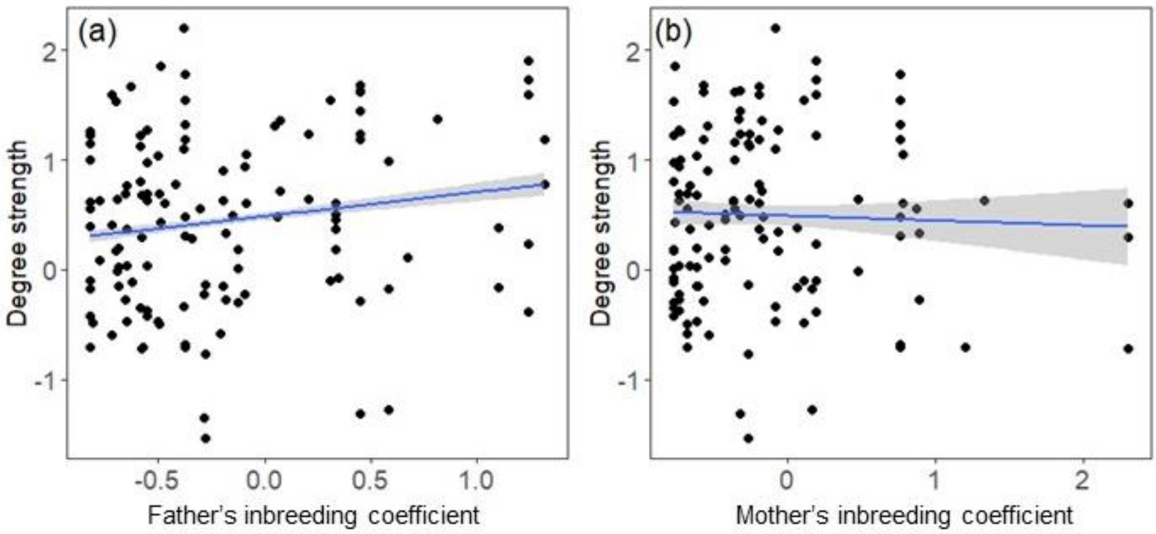


**Supplementary Figure 2**. Relationship between the weighted degree strength of each juvenile and their (a) father's inbreeding, excluding the two social fathers with extreme inbreeding coefficients, and (b) mother's level of inbreeding, excluding the two mothers with extreme inbreeding coefficients. Degree values and inbreeding coefficients are z-transformed. Line of best fit (blue line) with 95% C.I. calculated from model in Supplementary Table 3.

**Supplementary Table 1.** Characteristics of networks, relatedness, nest-box proximity and fledge date synchrony across the three study cohorts. Network characteristics describe the raw networks: thus, not all individuals may be included in analyses but they still contributed to the network characteristics of other individuals (e.g. number of associates). Average network association strengths are calculated from the raw network values and include all dyadic associations with strength > 0. Mean ± standard error values are given for network association strength, relatedness, nest-box distance, and fledge date synchrony; numbers in brackets for give the minimum and maximum values.

| Cohort | *N* juveniles | *N* network dyads with association = 0 | *N* network dyads with association > 0 | Network association strengths | Relatedness (*r*) | Inter-nest-box distance (km) | Fledge date synchrony (days) |
|---|---|---|---|---|---|---|---|
| 2015 | 33 | 354 | 246 | 0.11±0.006 | 0.07±0.003 (0.01 - 0.63) | 0.40±0.02 (0 – 1.54) | 22.40±0.67 (0 – 65) |
| 2016 | 75 | 2336 | 948 | 0.09±0.001 | 0.09±0.001 (0.02 - 0.70) | 0.62±0.04 (0 – 1.55) | 15.30±0.32 (0 – 106) |
| 2017 | 90 | 4838 | 1324 | 0.04±0.001 | 0.09±0.001 (0.02 - 0.70) | 0.60±0.01 (0 – 1.51) | 18.30±0.21 (0 – 89) |

**Supplementary Table 2.** Mean number of dyads for nests containing different combinations of sibling types, for each of the three cohorts.

| | Full sibling nest only | Half sibling nest only | Mixed-sibling nest |
|---|---|---|---|
| 2015 | 3.33 | 2 | 3 |
| 2016 | 3 | 2 | 4.5 |
| 2017 | 3.5 | 3.5 | 3.4 |

**Supplementary Table 3.** Results of linear mixed-effects models analysing the effect of inbreeding on network degree strength for three cohorts of juvenile hihi, with (a) two fathers, and (b) two mothers with extreme inbreeding coefficients removed from the analyses. P-values calculated using the double permutation method.

| | | Est. | S.E. | t-value | $P_{double\ permutation}$ |
|---|---|---|---|---|---|
| (a) | Intercept | 0.475 | 0.105 | 4.516 | |
| | Juvenile inbreeding coefficient (z-score) | 0.039 | 0.072 | 0.538 | 0.324 |
| | Mother's inbreeding coefficient (z-score) | -0.108 | 0.078 | -1.393 | 0.082 |
| | **Father's inbreeding coefficient (z-score)** | **0.272** | **0.164** | **1.662** | **0.036** |
| | Social father (yes/no) | 0.016 | 0.156 | 0.102 | 0.472 |
| | Father's inbreeding coefficient*genetic status (genetic/social father) | -0.172 | 0.273 | -0.632 | 0.107 |
| (b) | Intercept | 0.426 | 0.111 | 3.845 | |
| | Juvenile inbreeding coefficient (z-score) | 0.032 | 0.059 | 0.540 | 0.388 |
| | Mother's inbreeding coefficient (z-score) | -0.087 | 0.111 | -0.787 | 0.303 |
| | **Father's inbreeding coefficient (z-score)** | **0.139** | **0.074** | **1.871** | **0.017** |
| | Social father (yes/no) | 0.003 | 0.154 | 0.018 | 0.339 |
| | Father's inbreeding coefficient*genetic status (genetic/social father) | -0.104 | 0.161 | -0.645 | 0.258 |
